# Emergent regulation of ant foraging frequency through a computationally inexpensive forager movement rule

**DOI:** 10.1101/2022.02.15.480539

**Authors:** Lior Baltiansky, Guy Frankel, Ofer Feinerman

## Abstract

Ant colonies regulate foraging in response to their collective hunger, yet the mechanism behind this distributed regulation remains unclear. Previously, by imaging food flow within ant colonies we showed that the frequency of foraging events declines linearly with colony satiation ([1]). Our analysis implied that as a forager distributes food in the nest, two factors affect her decision to exit for another foraging trip: her current food load and its rate of change. Sensing these variables can be attributed to the forager’s individual cognitive ability. Here, new analyses of the foragers’ trajectories within the nest imply a different way to achieve the observed regulation. Instead of an explicit decision to exit, foragers merely tend toward the depth of the nest when their food load is high and toward the nest exit when it is low. Thus, the colony shapes the forager’s trajectory by controlling her unloading rate, while she senses only her current food load. Using an agent-based model and mathematical analysis, we show that this simple mechanism robustly yields emergent regulation of foraging frequency. These findings demonstrate how the embedding of individuals in physical space can reduce their cognitive demands without compromising their computational role in the group.

## 2 Introduction

Ant colonies rely on individual cognition and communication networks to perform complex collective tasks ([2]). Since brain tissue requires significant energetic investment ([3]), there is an advantage to communication systems that reduce the cognitive burden of the individual ([4]). It has recently been suggested that physical space can be utilised to offload computation from individuals’ cognition to their environment in the context of collective quorum sensing ([5]). This new principle may apply to other systems in collective behaviour, including foraging regulation.

Foraging in ant colonies is carried out by a small fraction of the workers, called foragers ([6]). When the foragers return to the nest, they distribute their harvest to other ants in the nest, and then re-exit to collect more food ([7]). This repetitive process persists as long as the food source is not exhausted or the colony satiates. Intriguingly, the rate at which food enters the colony matches the total level of hunger in the colony ([1], [8], [9]). The distributed nature of the ant colony dictates that this regulation emerges from local actions of individual ants.

Liquid food, such as honeydew or nectar, is commonly carried within an ant’s crop, an organ specialised for storing predigested food ([10]). From there it can be regurgitated to pass to other ants in mouth-to-mouth feeding interactions called *trophallaxis* ([11]). Trophallaxis is the main food-sharing method in many ant species ([12]). Each time a laden forager returns to the nest, she unloads the food from her crop to several receivers via trophallaxis. The food further circulates through a complex trophallactic network among all colony members ([9], [13]–[17]).

Previously, we used unique real-time measurements of fluorescent food inside the crops of all ants in a *Camponotus sanctus* colony, to infer quantitative links between local trophallaxis rules and the emergent regulation of food intake rate ([1]). The total level of hunger in the colony appeared to affect two aspects of the foragers’ behaviour. The first was the rate at which each forager unloaded her crop to receivers in the nest, which became slower as the colony satiated. Specifically, each forager’s unloading rate was proportional to the total “empty crop space” in the colony (hereinafter, ‘colony hunger’). The second was the average frequency at which each forager exited the nest for foraging. These individual foraging frequencies were, on average, linear with ‘colony hunger’. Our goal was to explore how these forager-colony relationships emerge from local rules.

The scaling of foragers’ unloading rate to total colony hunger was explained quite comprehensively by local trophallaxis rules that were identified from the empirical data ([1]). Progress has also been made toward revealing the local mechanism underlying the linearity of foraging frequency, though to a lesser extent. No precise immediate cause for a forager to exit the nest has been found. Contrary to past assumptions ([7], [8], [18]), foragers did not exit the nest only after they unloaded their entire crop contents, nor had we observed a clear crop-load threshold below which the foragers were more likely to exit. Rather, the foragers exited the nest with highly variable crop loads. Some studies have successfully identified local social triggers for the exits of foragers in several ant species ([19]-[26]), but most have focused on foraging initiation, and not on the subsequent decay of activity in response to gradual colony satiation ([27]). Our crop-load measurements in [1] have shed light on the local determinants of foragers’ exits, that linearly relate them to the current level of colony hunger. The local factors found to affect the temporal probability of a forager to exit the nest were both her instantaneous crop load, and her unloading rate in the nest: the emptier her crop and the faster her unloading, the more likely a forager was to exit. A simple Markovian decision-making model that yields the observed results was proposed. However, with no empirical access to the exact timings at which the forager assesses her next decision of whether to stay in the nest or leave to forage, the assumptions of our model could not be verified.

Here, we present an alternative mechanism, that reduces the cognitive demand on individual foragers through utilisation of physical space. This mechanism is supported by previously unexplored aspects of the data produced by our past experiments. The individual crop load dataset is now enriched with detailed spatial tracking of the foragers inside the nest. Together, these point to a new behavioural rule. A clear transition between two movement modes, depending on the forager’s instantaneous crop load, is evident from the new data: As foragers move around the nest, unloading their crops to ants that they meet, they tend to step towards the depth of the nest when their crop load is above a certain threshold, and tend to step towards the exit when it is below this threshold. Since the movement on both sides of the threshold is highly stochastic, this transition is masked when looking at the ultimate exit probabilities, which was the approach taken in [1]. Here we present an agent-based model that implements this new stochastic motion rule along with the previously reported trophallaxis rules, and analyze it mathematically. The results show that these rules suffice to shape a forager’s trajectory in the nest in such a way that produces linear foraging frequency regulation, while maintaining low levels of individual cognitive loads.

## 3 Results

### 3.1 Foragers move according to a biased random walk that is crop state dependent

Starved colonies of *Camponotus sanctus* ants were recorded as they gradually replenished on fluorescently-labeled food. All ants were tracked, the amount of food in their crop was quantified throughout time using fluorescence imaging, and all trophallaxis events were annotated ([1], [28]). Foragers were identified to be those ants that repeatedly left the nest to retrieve food and deliver it to other ants in the nest. We analyzed the trajectories of these foragers inside the nest in relation to their changing crop state, as they distributed their food in trophallactic interactions. Figure 1 shows a single frame from an experimental video overlaid with an example of a forager’s trajectory in the nest.

**Figure 1:**
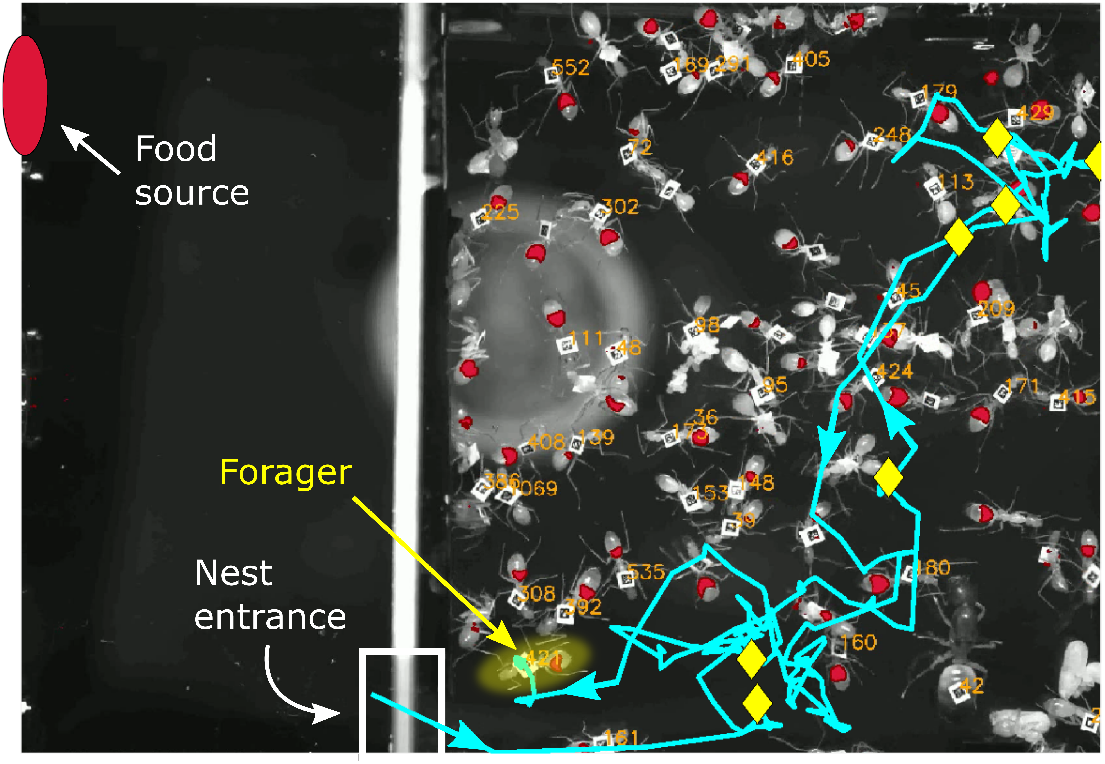
Example of a forager’s trajectory in the experimental nest. A single frame from an experimental video shows the nest on the right and a foraging arena on the left, where a fluorescent food source was presented. The food source is marked as a red oval and the nest entrance is marked by a white rectangle at the bottom left corner of the nest. Ant IDs are presented next to their tags, and the imaged food in their crops is overlaid in red. A single forager is highlighted in yellow, and her trajectory from when she last entered the nest is presented in cyan. Arrows on the trajectory mark the directionality of her path, and yellow diamonds mark locations of trophallactic interactions that she performed on her trip.

We found that the movement of a forager in the nest can be characterised by a random walk with a bias that depends on the amount of food in her crop (Figure 2A). At each trophallactic interaction that a forager performed, her distance from the nest entrance was measured. The probabilities of her next interaction to be farther from the entrance (step inward), closer to the entrance (step outward) or at the same distance from the entrance (stay), were calculated as a function of the forager’s crop state at the end of the interaction. Figure 2A shows that when the forager’s crop is more than 0.45 full, she is more inclined to step inward deeper into the nest. Conversely, at lower crop loads she is more probable to step outward toward the exit. Figure 2B shows that these probabilities are not affected by the direction of the forager’s previous step. Thus, it is reasonable to model the forager’s movement as a Markovian process.

**Figure 2:**
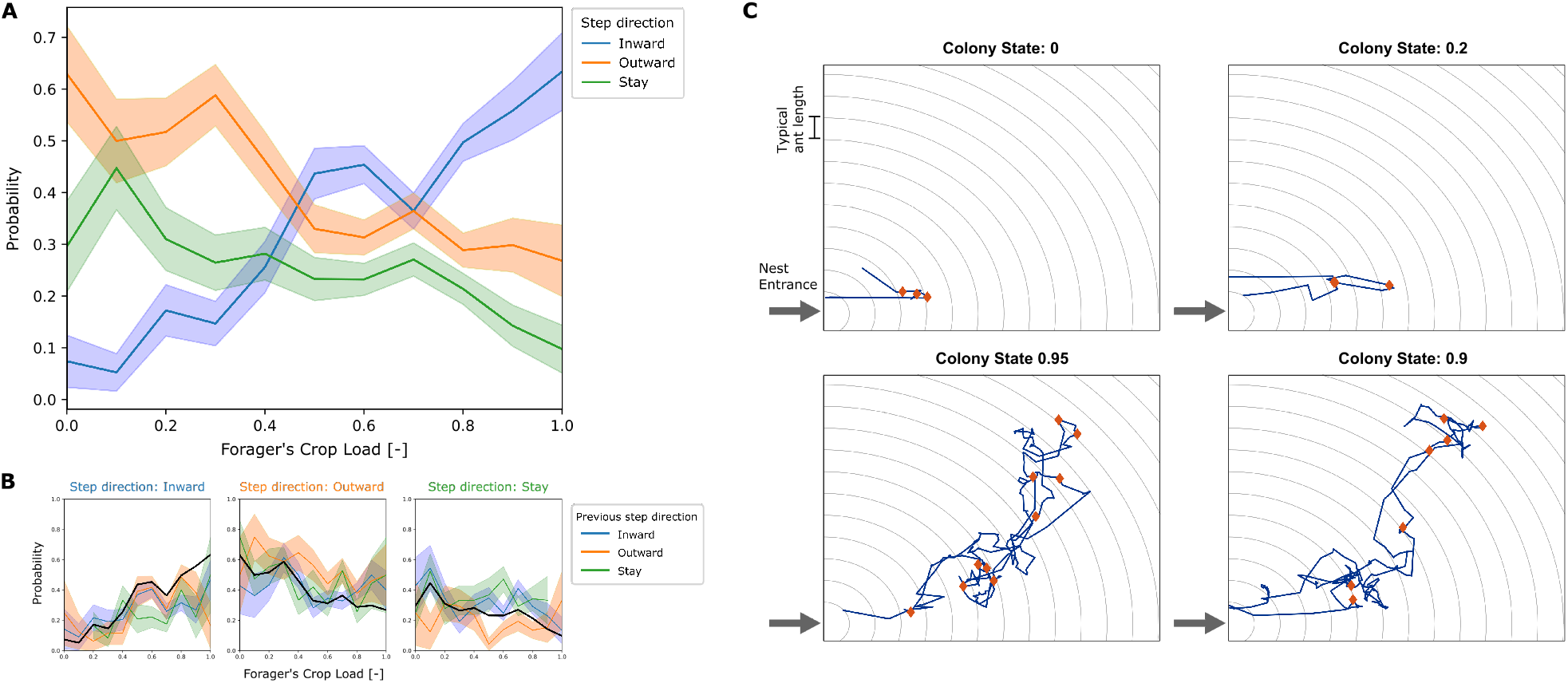
Empirical movement of foragers in the nest. **A.** All locations in the nest were binned according to distance from the entrance, with bin width of 1 typical ant length (as visualized by circular grid lines in panel C). At each interaction of a forager, her crop load at the end of the interaction and the location bin of her next interaction was recorded. Pooled data from all foragers was used to calculate the probability of the next interaction to be in a deeper location bin (inward), in the same location bin (stay), or in a location bin closer to the entrance (outward), as a function of their crop load. Probabilities and standard deviations were calculated for each one of 10 crop load bins. Standard deviation was calculated by the formula for multinomial 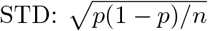, and is represented by the error bars in the plot. **B.** The data described in panel A was grouped according to the direction of the previous step. The plots show the probability to step inward (left), outward (middle) and stay (right), for cases where the previous step was inward, outward or stay as different curves. The pooled probability for all previous directions is presented as a thick black curve, equivalent to the curves presented in panel A. **C.** Examples of trajectories of single trips of a forager in the nest. Nest entrance is at the bottom left corner. Grid-lines spaced by a typical ant length are presented in gray. These are the spatial bins used to define the distance from the entrance for calculating the foragers’ biases (panels A-B). The trajectory of the trip is plotted in blue, and locations of trophallaxis events are presented as red diamonds. The top two plots present trajectories from low colony states, and the bottom two plots present trajectories from high colony states.

The crop-dependent movement described above could serve as a mechanism for generating random closed paths in the nest as foragers unload their crops via trophallaxis: since the forager steps into the nest with a relatively full crop after she fed at the food source, her initial bias drives her deeper into the nest. As she unloads her food to other ants, her crop load may reach a level at which her bias switches direction. The forager then continues to disseminate food to ants in the nest, but now with a drift that carries her toward the exit, until she finally reaches it and leaves the nest to forage again. Note that as the colony gradually satiates, the forager’s unloading rate decreases ([1]). Therefore the duration and the depth of the forager’s cyclic trips both rise with the colony’s level of satiety. Figure 2C shows examples of empirical paths of unloading trips of individual foragers in the nest. When the colony is hungry (colony state close to 0), the paths are short and include few trophallactic interactions, and when the colony approaches satiation (colony state close to 1), paths are long and include more trophallactic interactions.

To explore whether this empirically-derived movement rule may underlie the emergent linear relationship between foraging frequency and total colony hunger, we simulated it numerically and analysed it mathematically. In the next sections we present two agent-based models and an analytical description of the system. The first agent-based model mimics the experimental 2-dimensional nest. The second model is a simplified 1-dimensional version which is more readily approachable analytically. Our simulations show that both models yield the desired linear foraging frequency regulation. We then solve the 1D model analytically to show how it accounts for this emergence based on the local movement rule described above. Finally, we compare different properties of these models to our empirical observations.

### 3.2 Agent-based model in a 2D nest

This model implements a 2D nest consisting of 81 cells (9×9), each containing one ant. The size of the nest and the number of ants were chosen to be of similar scale to the experimental conditions. The nest has a single point of entry/exit, located at one of the corner cells, mimicking the structure of the experimental nest. A single forager moves according to the crop-dependent statistics described in section 3.1, bringing food into the nest and distributing it among the inhabiting ants. For simplicity, the non-forager ants only receive food from the forager and do not redistribute it further. Note that contrary to the assumptions used in our previous paper ([1]), here a forager never decides to exit the nest. Rather, an exit occurs if the forager’s motion brings her to the nest exit. Foragers that exit the nest are modelled to refill their crop and then re-enter the nest. Hereafter, we refer to all the steps between the forager’s entrance and exit as a single *trip*. For more details please refer to the Materials and Methods section.

The simulation implements three simple rules that were derived from the experimental data.

1. **Forager movement.** The forager moves according to the biased random walk described above, choosing a random cell of its 8 neighbours to move to, with probabilities derived from the empirical data (Table 1).
2. **Trophallaxis.** At every step, the forager performs trophallaxis with the nest-ant that shares her current position. The amount of food passed from the forager to the nest-ant is stochastic, and is scaled to the available crop space of the receiver ant. As observed empirically, it is a random, exponentially distributed, fraction of the receiver’s unfilled crop space, with an average of 0.15 ([1]). If the forager has insufficient food, she gives all that she has.
3. **Nest-ant movement.** Nest-ant movement is implemented in the model between each trip of the forager: each time the forager exits the nest, each nest ant moves up to 4 cells in a random direction. This quantity was chosen based on empirical data, which shows that nest-ants moved a mean ± STD of 2.83 ± 1.45 ant-lengths between consecutive foraging trips. Nest-ant movement contributes to the spatial homogenization of the food in the colony, which causes the forager to interact with ants that are, on average, representative of the satiety state of the whole colony (see section 3.5). This unbiased sampling was observed empirically, and together with the empirical trophallaxis rule, causes the forager to unload her crop at a rate proportional to the colony’s total hunger level ([1]).

**Table 1:** Movement biases for agent-based model. The probabilities of a simulated forager to step inward, outward or to stay in the same space bin, for two cases: when her crop load is lower than or higher than a threshold (0.45). The values of the threshold and the biases are approximated based on the empirical data (Figure 2A).

| Crop load | $P(inward)$ | $P(outward)$ | $P(stay)$ |
| --- | --- | --- | --- |
| $\leq 0.45$ | 0.16 | 0.53 | 0.31 |
| $> 0.45$ | 0.46 | 0.32 | 0.22 |

**Table 2:** Parameter values for different groups of ants. Parameters given to all agents are described under the ‘Ants’ sub-population.

| *Sub-population* | *Parameter* | *Model* | *Value* |
| --- | --- | --- | --- |
| Ants |  |  |  |
|  | Crop state capacity | - | 1 |
| Non-foragers |  |  |  |
|  | Initial crop state | - | 0 |
|  | Position |  |  |
|  |  | 1D | 45 ants, one on every cell |
|  |  | 2D | 89 ants, one on every cell |
| Foragers |  |  |  |
|  | Initial crop state | - | 1 |
|  | Threshold value | - | 0.45 |
|  | Initial position | - | Entrance of nest |
|  | Biases in state A | - | {0.32, 0.22, 0.46} |
|  | Biases in state B | - | {0.53, 0.31, 0.16} |
| | Interaction proportion | - | $p \sim \text{Exp}(\frac{1}{0.15})$ |
|  | Interaction rate | - | Every step |
|  | Foraging time | - | 0 |

The simulation qualitatively reproduced the lengthening and deepening of foragers’ trips with colony satiation that were observed empirically (compare Figure 3 and Figure 2C). Figure 3 depicts two trips of the forager within the simulated nest, from different stages of a single run of the simulation: one from an early stage of the run, when the colony was 5% satiated, and the second from a later stage, when the colony was 90% satiated. These representative examples demonstrate how the same set of unloading and movement rules by which the forager operates, produces short trajectories when the colony is relatively hungry, and longer trajectories as the colony satiates.

**Figure 3:**
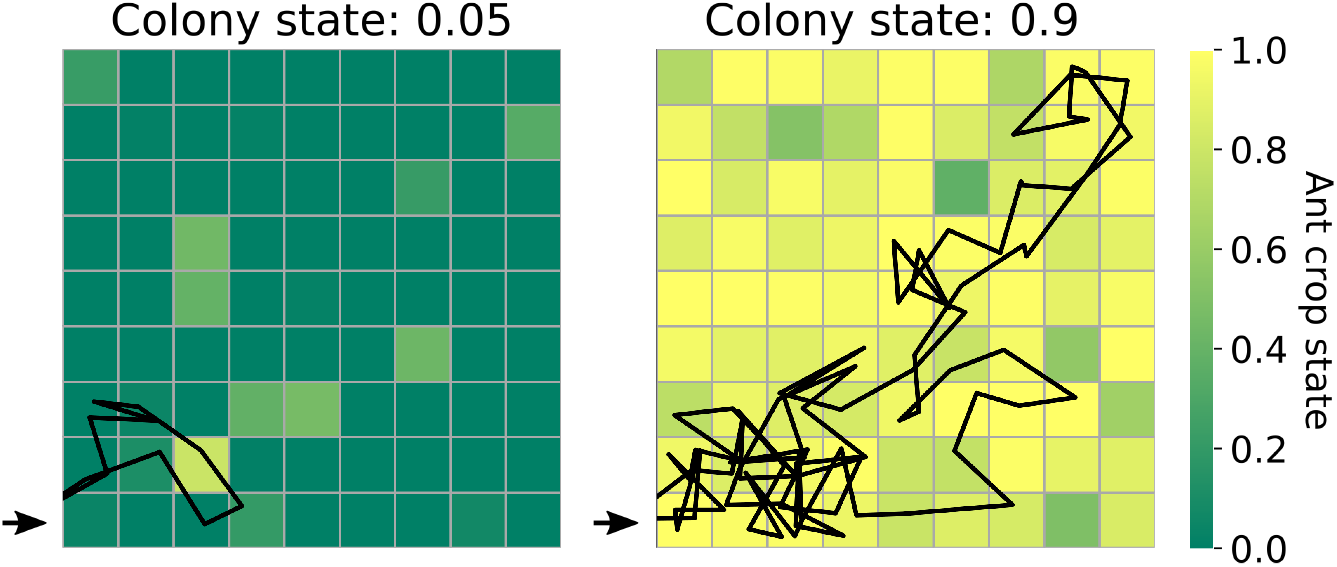
Examples of two trips of a forager in the 2D simulation. The nest is presented as a 9×9 grid, in which the colour of each cell represents the crop state of the ant that inhabits it (dark green represents an empty crop, yellow represents a full crop). The nest entrance is marked by a black arrow at the bottom left corner of the nest. All of the forager’s positions during the trip are presented as a black trajectory through the cells. (For the sake of visualisation, a small random component was added to each position to avoid overlaps in the plot whenever the forager revisits a cell.)

Moreover, the dynamics of the lengthening of the trajectories in the nest indeed lead to a linear relationship between the average frequency at which the forager encounters the nest exit, and the amount of food accumulated in the colony. This is analogous to the linear matching of foraging frequency to colony hunger that was observed empirically (Figure 4A and B).

**Figure 4:**
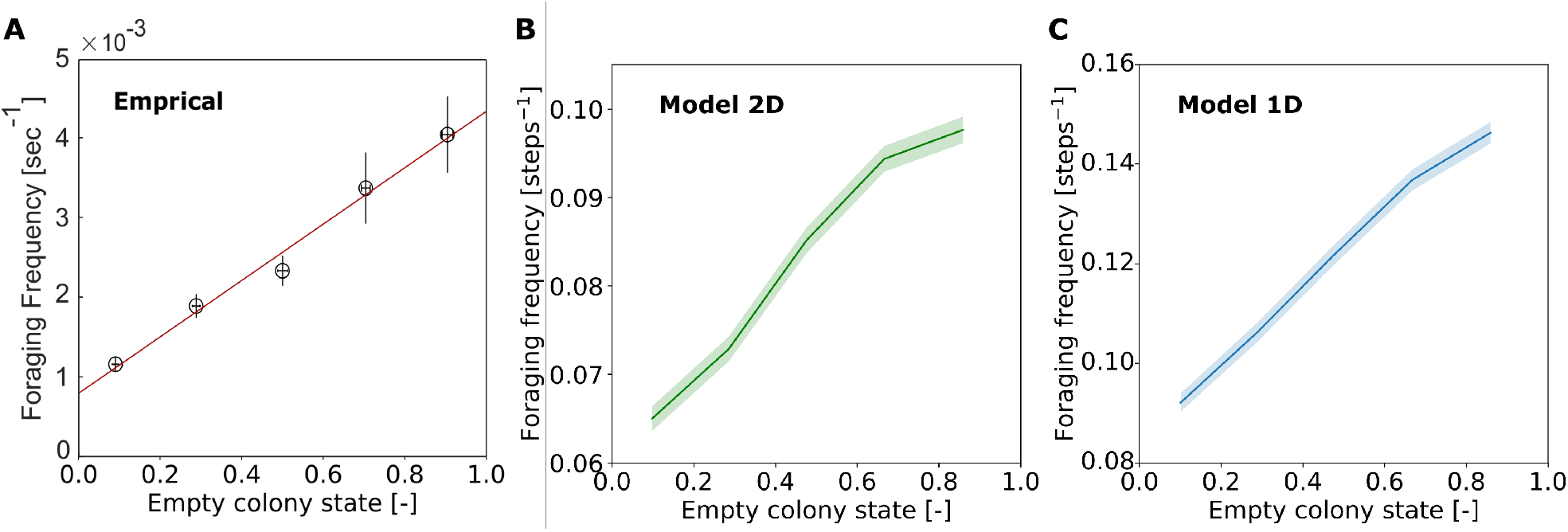
Linearity of foraging frequency with empty colony state. Foraging frequency was calculated as the inverse of the duration of the forager’s trip in the nest. Trips were binned into 5 equally-spaced bins of colony state, and the mean and SEM of foraging frequency was calculated for each bin. **A.** Experimental data, figure taken from [1]. **B.** Data from 200 repeats of the 2D model simulation. **C.** Data from 200 repeats of the 1D model simulation.

Hence, the empirical foraging frequency regulation on the colony level emerges from the 3 local rules implemented in our agent-based model. Since the direction of the forager’s movement in the nest is coupled to the amount of food that she carries, and given that her rate of unloading is determined by the amount of food in the receivers’ crops, there emerges a negative feedback between the amount of food stored in the colony and the frequency at which foragers exit the nest to bring in more food. This cross-scale feedback, from the level of colony hunger to the level of individual foraging events, emerges with no need for the forager to sense anything but her own current crop load.

To understand how this linearity emerges from the local rules described above, we first simplify the system to one dimension, which is more readily approachable analytically. Since the direction of the forager’s movement is defined relative to the nest entrance (toward the entrance, away from the entrance), the forager’s position may essentially be defined with one coordinate - her distance from the entrance. The nest can then be simplified to a 1-dimensional array of nest-ants through which the forager walks back and forth. In the next section we describe the 1D version of our agent-based model.

### 3.3 Agent-based model in a 1D nest

This model implements a 1D nest consisting of 45 cells, each cell inhabiting one nest-ant. The point of entrance/exit of the nest is from one of its edges. A single forager walks in the nest and feeds nest-ants as described in the 2D model above, with the following adjustment to the nest-ant movement implementation: Instead of each nest-ant moving up to 4 cells in a random direction, all nest-ant positions are randomly shuffled between forager trips. This adjustment is required for sufficient homogenization to yield a representative sample of receivers for the forager, and is supported by 1-dimensional projection of empirical nest-ant data (see details in SI). For more details on the implementation of the 1D model, see Materials and Methods.

Figure 4C shows that, similar to the 2D model, the 1D model reproduces the emergent linear relationship between foraging frequency and colony hunger as was observed empirically. Next, we present a mathematical description of the 1D system to analytically explain these results.

### 3.4 Emergent linear relationship between foraging frequency and total colony hunger

A precise analytical description of the dissemination trips of a forager is challenging, since they involve stochasticity in her movement, in the amount of food she delivers at each interaction, and in the state of her nest-ant partners. Therefore, we use a coarse-grained analysis, where we consider the averages of these stochastic variables: the forager’s average direction, the average amount of food given per interaction, and the average state of the forager’s partners in a trip.

Here, for the sake of simplicity, we present the equations for the extreme, deterministic case in which a forager walks only inward when her crop load exceeds the threshold, and only outward when it is below the threshold. In the Supplementary Information we show how these equations apply to the more general case, where the forager’s bias is set with partial probabilities to walk in each direction.

Let *c* denote the crop state of the forager (*c* =1 when the forager’s crop is full and *c* = 0 when it is empty). A certain crop load *c*\* is the threshold that separates between the forager’s two movement biases within the nest: the forager walks inward when *c* > *c*\* and outward when *c* < *c*\*.

Let *F* be the total satiety state of the colony (*F* = 0 when the colony is starved and *F* = 1 when all ants in the colony are satiated). The nest-ant movement rule dictates that the forager interacts with a representative sample of the colony at each trip ([1]), such that the average state of the forager’s partners is equal to the colony state, *F*. The trophallaxis rule gives the average amount of food delivered at each interaction: a fraction *α* of the receivers’ empty crop space ([1]). Given these two rules, the average amount of food a forager unloads at every interaction is:

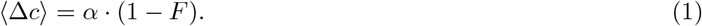

Since the forager enters the nest full, and since in the extreme case the forager performs trophallaxis with a new ant at each step, the average number of interactions she will make until her crop load reaches the threshold is 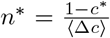. In the extreme case, this quantity is equivalent to the average position in the nest at which the forager switches her bias, denoted 〈*x_switch_*〉. Therefore, we obtain the following relation between the colony state and average position at which the forager switches her bias:

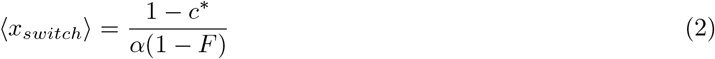

The average duration of the foragers’ trip, 〈*T*〉 is the time it takes her to reach 〈*x_switch_*〉 from the entrance (at *x* = 0) and return. In the extreme case, walking inward every step until 〈*x_switch_*〉 and outward every step from 〈*x_switch_*〉, this simply equals 2 · 〈*x_switch_*〉. We get:

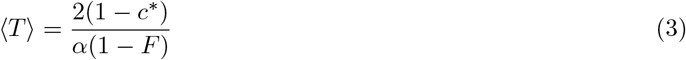

The frequency of the forager’s trips, *R*, is defined as the reciprocal of the average trip duration 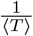. Therefore, the foraging frequency is:

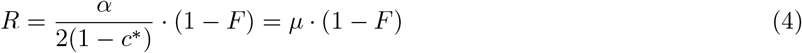

where 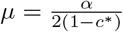 is a constant. Thus it is clear that the foraging frequency *R* is proportional to the colony state of hunger (1 – *F*). This is the linear relationship observed both experimentally and in the simulations of our agent-based models (Fig. 4).

Clearly, in reality forager ants don’t move in such an extreme manner within the nest, but the general logic of the analytical development above applies to a softer movement rule as well, where the forager’s walk is more probabilistic. In short, the difference between an extreme walk and a probabilitic walk, means that the forager, going stochastically back and forth between the nest ants, may interact multiple times with same ants before switching her bias. This distinction alters the average amount of food delivered at each step (Δ*c*, eq. 1), as less food is given to a nest-ant with each subsequent encounter between her and the forager. Additionally, the number of interactions that it takes the forager to reach her threshold no longer translates directly to the position at which she switches her bias (*x_switch_*, eq. 2). Nevertheless, it turns out that the average amount of food given to each nest-ant is still proportional to (1 – *F*), and that since both the inward and outward biases are constant, the number of steps spent with each nest ant is, on average, also constant (neglecting boundary effects). Therefore, overall, the differences introduced by the probabilistic walk are expressed in the factor *μ* that multiplies the colony state of hunger (1 – *F*) in equation 4. In the probabilistic movement case, *μ* is dependent on the fraction *α*, the threshold *c*\*, and the probabilistic walking biases. For details, see SI.

### 3.5 Further trip characteristics in experiment and simulation

Figure 5 presents additional dynamics that appeared in both the experimental and simulated data. The states of the foragers’ recipients represent, on average, the states of all ants in the colony (Fig. 5A). The forager unloads her food at a rate proportional to the empty colony state (Fig. 5B). This is a direct result of the trophallaxis rule (volume proportional to recipient empty crop space), the constant trophallaxis rate (trophallaxis at every encounter with a nest-ant), and the nest-ant movement rule that yields representative receivers (Fig. 5A). Furthermore, the forager’s trips in the nest become deeper with colony satiation (Fig. 5C), and the crop states with which the forager exits the nest are highly variable at all colony states (Fig. 5D). On average, they are relatively constant initially, and slightly rise at higher colony states. The trip depth and exiting crop dynamics are more similar to the empirical data in the 2D version of the model.

**Figure 5:**
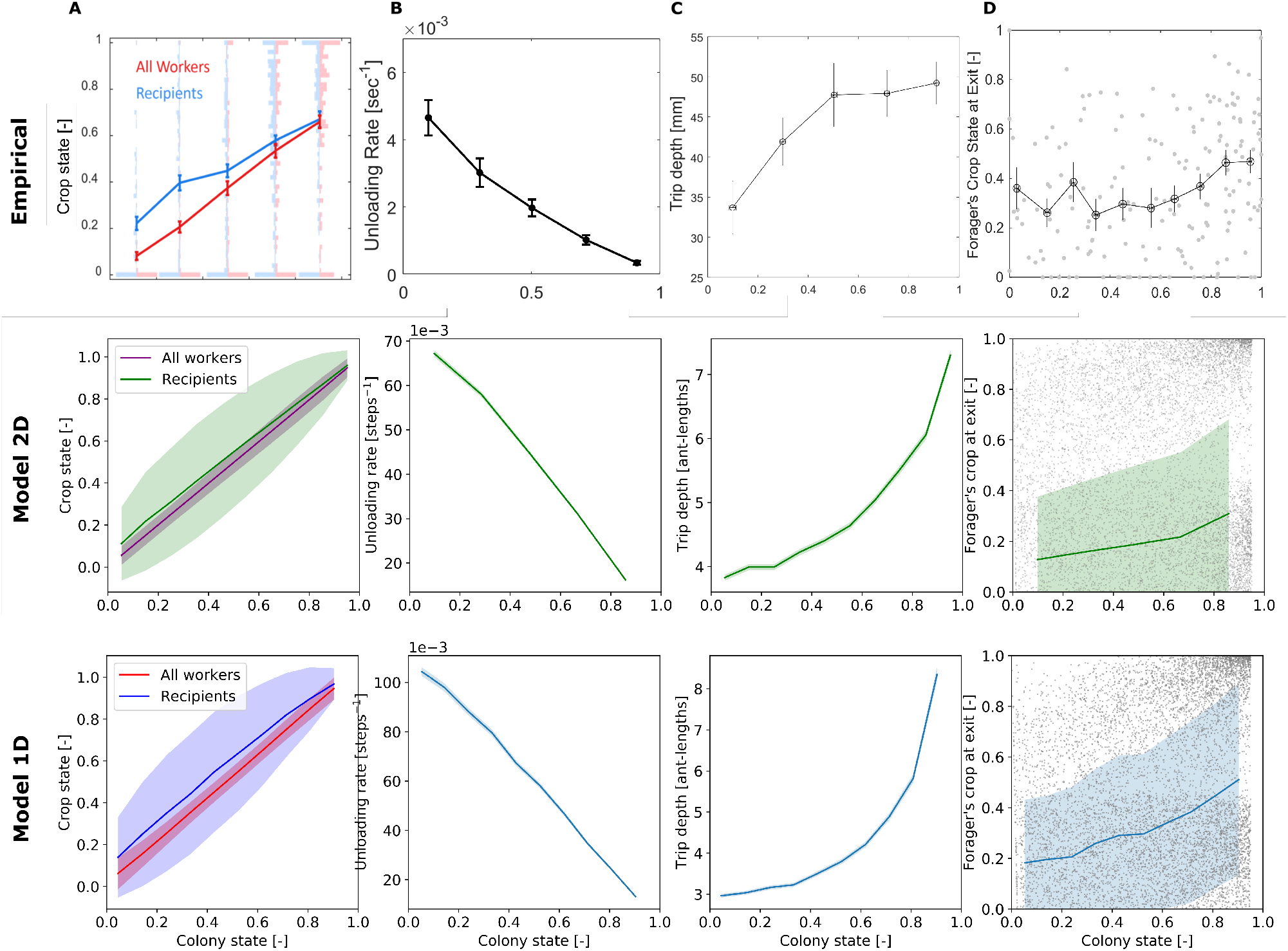
Comparison between empirical and simulation data of forager trip dynamics. Trips were binned into equally-spaced bins of colony state. Means and SEMs of different measures were calculated for each bin. Plots of the empirical data were taken from [1]. **A.** Average crop state of the foragers’ trophallactic partners in the trip and of all nest-ants. Error bars for the simulated data represent STDs to better appreciate the variance of the distributions. **B.** The forager’s unloading rate, calculated as the amount of food she delivered in the trip divided by the duration of the trip. **C.** Trip depth, calculated as the maximal distance of the forager from the nest entrance in the trip. **D.** Forager’s crop load at the end of the trip. Error bars for the simulated data represent STDs to better appreciate the variance of the distributions.

Although the trend of increasing trip depths was captured in both of our models (Fig. 5c), there is a qualitative difference between the shapes of the empirical and simulated curves. While the empirical rise in trip depth is concave and seems to reach a plateau, the resulting curves of the agent-based models are convex (especially that of the 1D model). We speculate that three features of the empirical system that are not incorporated into the agent-based models may be the cause for this minor inconsistency. The first is that in reality ants occupy space in the nest, thus restricting the movement of other ants by steric interactions. That is, the forager’s state space may be constrained since she may be blocked from reaching certain areas of the nest by other ants. On the other hand, in the model, the forager is able to walk over nest ants and hence has no state space restriction. The second feature is that the model does not implement trophallaxis between nest ants, whereas it is known that in real ant colonies nest ants do indeed spread food between themselves. Empirically, this nest ant behaviour may be a reason that the forager does not need to cover all areas of the nest and may be a reason for the plateau in Figure 5 Empirical C. Lastly, the third feature is the spatial distribution of ants in the nest. While in the models nest-ants occupy all cells in the nest, the empirical distribution of ants is usually characterised by dense regions of less mobile ants and sparse regions where ants tend to move more. Naturally, foragers’ interactions with nest-ants may occur only where nest-ants are present, thus affecting the locations where foragers are found in the nest.

## 4 Discussion

Ant colonies manage to regulate foraging activity in response to their collective hunger, despite the fact that the foragers are only a small subset of the workers. In [1] we have shown that as starved colonies gradually satiate, the average individual foraging frequency linearly matches the temporal state of hunger of the whole colony. Here we have presented new experimental data that accounts for this relation between the colony scale and the individual scale. Combining spatial tracking of foragers within the nest with the dynamic measurements of their crop loads, our data implies a simple rule for the movement of foragers, that ties their instantaneous crop load to the direction that they take within the nest. Overall, our findings suggest that a forager’s trajectory in the nest can be shaped by the rate at which she unloads to recipient ants, while this unloading rate is governed by the satiety of the recipients. Using an agent based model and mathematical analysis, we demonstrated that together with the trophallaxis rules described in [1], this simple movement rule produces linear foraging frequencies as observed empirically.

Previous studies have identified local factors that may determine foraging activity in various species of social insects. These factors include chemical cues and the rate of interactions with other individuals ([19]-[24], [29], [30]), the foragers’ own nutritional state ([31], [32]), and larval hunger signals in the nest ([33]-[43]). Such factors were shown to relate the foragers’ activity to external variables such as food quality or availability, and to the internal colony nutritional requirements ([44], [45]). However, no study that we know of provides a full mechanistic explanation for the qualitative linear relationship between colony hunger and individual foraging frequency, which was observed during the process of gradual colony satiation. This striking linearity was revealed only recently, thanks to technological advances ([1]).

In our previous work ([1]), we have presented a model which described the forager’s decision to exit as a function of both her crop load and her unloading rate, however it did not present a comprehensive mechanism of action. In the current study, we show that the exiting rate dynamics can be described without an explicit decision to exit by the forager, rather, the forager only decides on the direction of her next step, leaving the nest to forage whenever she reaches the nest exit. Indeed, empirical data supports that such decisions are Markovian and depend on a single variable - the forager’s current crop load. In comparison to the previous model, this model alleviates the forager’s need to keep track of her unloading rate. Sensing the unloading rate, that is the change in crop load over time, requires some form of memory of past crop load values. Therefore, the new model presents a simpler mechanism by removing this computational and memory burden.

Other than its simplicity, this model is preferable over the previous one for its greater explanatory power. It manages to explain both the linear foraging frequency and the deepening of foragers’ paths in the nest, implying that both of these trends result from the same set of rules. The deepening of foragers’ visits with colony satiation may be a widespread phenomenon, as it was also observed in honeybees ([44]).

Additionally, our analytical understanding of the described behavioural rules emphasises the robustness of the system to intrinsic forager parameters, such as threshold value and bias strengths. So long as there is a crop load at which the forager’s movement bias switches from inward to outward, her exiting frequency is expected to be linear with her unloading rate. Since the forager’s unloading rate is controlled by her recipients, her exit frequency is linked to the colony. This allows for different foragers to have different movement biases even within the same nest, and still the relationship between their exit frequency and colony satiation will remain intact.

On the other hand, the sensitivity of the forager’s unloading rate to the crop loads of the ants she encounters, means that a linear relationship between her foraging frequency and the total colony hunger requires her receivers to be representative of the colony. While our experiments indeed display a representative sample of receivers and a linear relationship with colony state, our model predicts that different interaction patterns will yield different results. In cases where the forager encounters non-representative subgroups of the colony, her foraging frequency is expected to be linear with the state of her sample, but this may no longer translate to linearity with the collective state of the colony. In nature, ants’ nests are typically composed of multiple chambers ([46]-[48]), thus the nest-ant distribution is more clustered and organized than in the artificial single-chamber nest that was used in our experiments ([49]). Accordingly, it may be that nest architecture will affect the sampling characteristics of the foragers, and consequently their foraging frequency ([50], [51]). Other factors that may affect the forager’s sample include the number of nest entrances ([52], [53]), the density of ants in the nest, and the topology of the colony’s trophallactic network ([9], [14]–[17], [54]–[57]). Fortunately, modern tracking methods enable to acquire more data on trophallactic networks to explore these potential effects ([28], [58]).

In a broader view, the notion that the movement rule described in this paper relieves the forager’s need to keep track of past measurements highlights a unique property of the role of space in a system. By nature, an animal’s position in space is the integration of all of its movements. Therefore, a system of agents that translate a certain variable into movement in space, naturally encodes aspects of the history of this variable as well, with no need for the agents themselves to allocate memory and perform computations on this information. This principle can be useful for distributed systems composed of simple computing elements ([59]): memory in the system emerges simply from being embedded in space ([60]).

Recently, Pavlic et al. have demonstrated a similar externalisation of computation onto physical space in a multi-agent system ([5]). They offered a new mechanism for quorum sensing, a ubiquitous phenomenon in animal groups and a useful process in distributed systems. While previous algorithms for quorum sensing relied either on the communication between agents or on the ability of agents to accumulate information on their encounters with other agents, the mechanism introduced by Pavlic et al. utilised physical space to relieve these computational demands from the agents. They reasoned that since accumulating information in an animal’s brain is believed to rely on neuronal dynamics that are analogous to a random walk ([61]-[63]), when a decision is made by a group of animals, this neuronal random walk can be instantiated in the physical movement of the individuals of the group instead. Thus, computation is externalised from the individuals’ brains to the physical environment in which they move. A similar mechanism, wherein a cognitive decision variable is encoded in the mutual location of an entire group was recently presented for *P. longicornis* ants ([64]).

Similarly, the model we present here for foraging frequency regulation supports the same notion. If the movement of the foragers is neglected, it may seem that in order for foraging frequency regulation to emerge, foragers must accumulate information on their changing crop load in order to decide when to exit the nest ([1]). However, here we show that once forager movement is considered, the decision when to exit the nest is no longer an internal decision of the forager, but an external decision made by the collective that includes the forager, the colony, and their physical environment. Accordingly, the internal behaviour of the forager then solely relies on her current crop load, with no need for her to accumulate information on its history. The cumulative information on past crop load values is represented by the forager’s position in the nest, and is thus stored externally in physical space. This exemplifies how utilisation of an individual’s position in space can reduce its cognitive demands without detracting from its computational contribution to group-level emergence.

## 5 Materials and Methods

### 5.1 Experimental setup

The experiments used to conduct this research are those used in [1].

### 5.2 Data analysis

Data was analysed in Python using the following packages: Numpy ([65]), Matplotlib ([66]), openCV ([67]) and Pandas ([68]).

### 5.3 Agent-based model

The agent-based models are described according to the protocol laid out by Volker Grimm, et al. ([69], [70]). The model includes a single forager which has two walking tendencies, in either a 1-dimensional or 2-dimensional nest. When she has food in her crop, such that she is above a threshold value, she tends deeper into the nest. Once her crop level drops below the threshold, she tends towards the entrance. The forager begins at the entrance with a full crop and walks through the nest, unloading food to each non-forager she meets according to her trophallaxis rule. Once she has unloaded enough food, she switches her walking tendency, and walks towards the entrance. Upon reaching the entrance, she refills and proceeds to re-enter the nest.

The model is written in Python with a GUI written in Java. Scheduling was carried out through a modified version of the mesa scheduling module ([71]).

#### 5.3.1 Purpose and Patterns

The purpose of the model is to determine whether the three rules described in section 3.2 are sufficient to recapitulate forager trip frequency relation with colony hunger. Size of the nest, nest coverage and other parameters not used in this paper are designated prior to running of the simulation. Global and other forager patterns are used to determine the accuracy of the model. Such dynamics include forager depth per trip, forager exiting crop, unloading rate, among others.

#### 5.3.2 State variables and Scale

The model is comprised of individual agents representing ants; ants can be grouped into two sub-populations, foragers and non-foragers. These two populations are representative of what is seen in ant colonies in the scope of food dissemination. The model is also treated as an individual object to allow for data collection and parameter setting.

Biases {a, b, c} are to be read as such; a is probability to step backwards, b is probability to stay, c is probability to step forward.

In both models one forager was initialised and simulations were run until the colony was sufficiently satiated. Nest are made up of cells which are taken to be the size of a single ant. The models differ in their size and nest ant deployment.

##### 1D model

The nest length is 45 cells, plus 1 entrance cell. The entrance and deepest cell in the nest are reflecting boundaries, forcing the forager to step forwards/backwards, respectively, the step after it reaches said cell.

##### 2D Model

The nest dimensions is 9 by 9 cells, plus 1 entrance cell.

#### 5.3.3 Process overview and scheduling

The process of the forager in all models is described by the flow diagram in figure 6.

**Figure 6:**
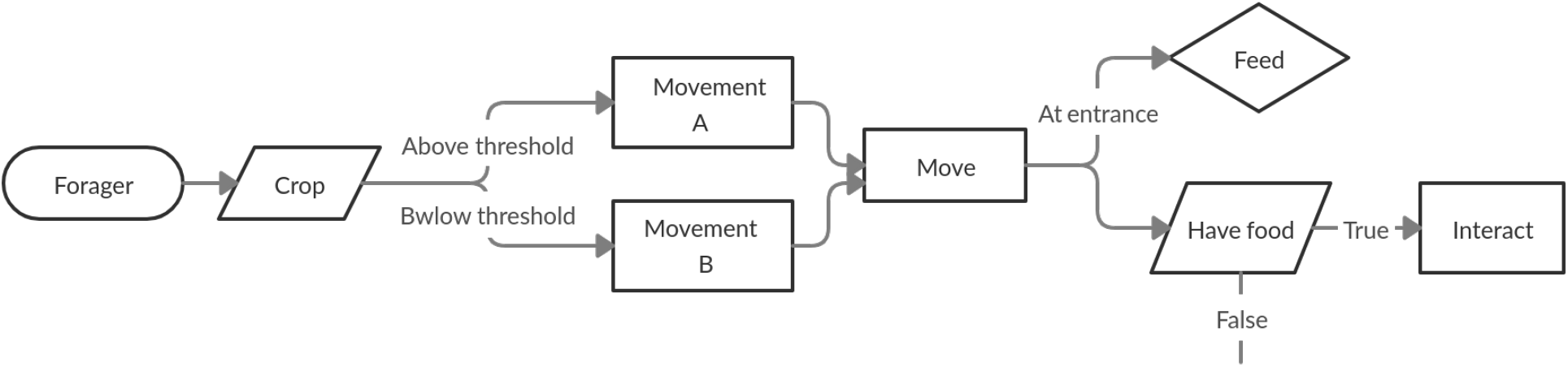
Schematic detailing the process of a forager at every step of the simulation. The forager first moves according to the bias random walk (defined by its crop). Then either feeds, if at the entrance, or attempts to interact with another agent.

Time in the model is discrete, with each step in the simulation representing the time it takes for the forager to step one ant length. Although there is a single forager in the model used for these results; the model is written with asynchronous, sequential scheduling as more foragers can be added. Random scheduling was applied as each forager is assumed to not be actively cognisant of other foragers’ actions, hence act independently.

#### 5.3.4 Design concepts

##### Emergence

Forager dynamics emerge from the behaviours of the model. Interactions and movement are hard-coded, however, dynamics such as trip duration/depth, exiting crop and colony state progression all develop only as a consequence of these behaviours. Hence, the foragers’ adaptation to the changing colony state occurs implicitly via these behaviours and its position.

##### Sensing

All agents are assumed to know their crop levels, and on top of this, the forager also knows a movement threshold crop level. Agents are not assumed to know where they are in the nest or any information about other agents/the system.

##### Interactions

Trophallaxis between agents is modelled explicitly in the trophallaxis rules.

##### Stochasticity

Trophallaxis and movement of foragers are both modelled as stochastic behaviours.

##### Observation

Simulations were repeated a number of times, with crop state of all agents at every step averaged over these repeats. Forager specific data; crop, position, current trip length, was recorded at every step for every individual run of the simulation. Interactions volumes and partners were recorded for every step in every individual run of the simulation. Trip duration and exiting crop were recorded every time the forager returned to the entrance for every individual run of the simulation.

#### 5.3.5 Initialisation and termination

Every simulation was initialised with empty non-foragers at every position in the nest and a fed forager at the entrance. This mimics the data collected from wet-lab experiments, in which colonies were starved for 1-2 weeks prior and data collection only began after the first time a forager leaves the nest to find food.

Simulations were terminated after all nest-ants were at least 95% full (4000 steps for the 2D model and 2000 steps for the 1D model).

#### 5.3.6 Input

No external input into the models was used.

#### 5.3.7 Sub-models

##### Forager movement

At every step, the forager can move deeper into the nest, move towards the entrance or stay in the same position. This decision is a probabilistic process defined by walking biases that were extracted from empirical data. The biases differ depending on whether the forager’s crop is above or a below a threshold value of 0.45 (also extracted from empirical data). Based on the threshold, the forager can be in two states; state *A* in which she is tends deeper into the nest and state *B* in which she tends towards the entrance. The transition between these states occurs when her crop falls below the threshold.

##### 1D Non-forager movement

Every time the forager returns to the entrance, she feeds and the order of the ants in the nest is randomly shuffled, after which, the forager begins its next visit into the nest. Random shuffling of non-foragers is taken from empirical data as discussed in the representative sample rule in section 3.2.2.

##### 2D Non-forager movement

Every time the forager returns to the entrance, she feeds and nest ants are able to move to any position within their Moore-neighbourhood with a radius of 4.

##### Trophallaxis

At each step, the forager checks whether it shares a cell with a non-forager, in this model, this is always true unless the forager is at the exit. If there is a non-forager, the forager is not empty and the non-forager is not full, the two agents perform unidirectional trophallaxis, from forager to non-forager. The forager transfers a proportion of the recipients empty crop space. The proportion is sampled from an exponential distribution with a mean of 0.15. The amount transferred from forager to recipient is called the interaction volume.

## 6 Acknowledgements

We thank Efrat Greenwald for collecting the empirical data used in this study. Thanks to Amos Korman and Jean-Pierre Eckmann for mathematical consultation and ideas, and to Aviram Gellblum for coding advice. This research was supported by the Minerva Foundation, the Israel Science Foundation, Grant No. 1727/20, the European Research Council (ERC) under the European Unions Horizon 2020 research and innovation program (Grant agreements No. 648032 and 770964).

## 7 Competing interests

The authors declare no competing interests.

## 8 Data availability

- **Figure 2 - source data 1: Empirical data.** Legend: Data used for calculation of the foragers’ empirical crop-dependent bias. All foragers’ interactions are pooled from the 3 experiments presented in [1]. Each interaction entry includes information on its location in the nest, the direction of the next interaction of the forager, and the forager’s crop load.
- **Figure 4 - source data 1: Data from 1D model.** Legend: Output data from 200 runs of the 1D agent-based model. The file contains 3 spreadsheets: (1) Forager data. Includes data on the forager’s crop load and position in the nest at every step of the simulation. (2) Trophallaxis data. Includes data on the forager’s and the receiver’s crop loads, and the amount of food transferred at every interaction. (3) Trip data. Aggregated data on each trip of the forager inside the nest, including trip length and forager’s crop load upon exiting.
- **Figure 4 - source data 2: Data from 2D model.** Legend: Output data from 200 runs of the 2D agent-based model. Data within the file is as described for the 1D model data.
- Python code for the agent-based model is available on GitHub ([72]).

## Supplementary Information

### Nest-ant movement in 1D model

To translate the empirical 2D movement of nest-ants to 1D for the 1D agent-based model, we mapped all nest-ants’ 2D positions into a 1D ranking according to their euclidean distance from the entrance. To quantify the degree of their movement between consecutive foraging trips, we calculated the Kendall *τ* rank correlation index for each pair of consecutive sets of rankings. The coefficients for each one of our 3 experiments were distributed close to 0, indicating that the change in ants’ rankings was close to what would be obtained by random shuffling. Indeed, we also randomly shuffled the empirical rankings for comparison, and the resulting coefficients were not different from those calculated on the non-shuffled empirical rankings (Figure S1). Therefore, nest-ant movement in the 1D model was implemented as random shuffling between consecutive foraging trips.

**Figure S1:**
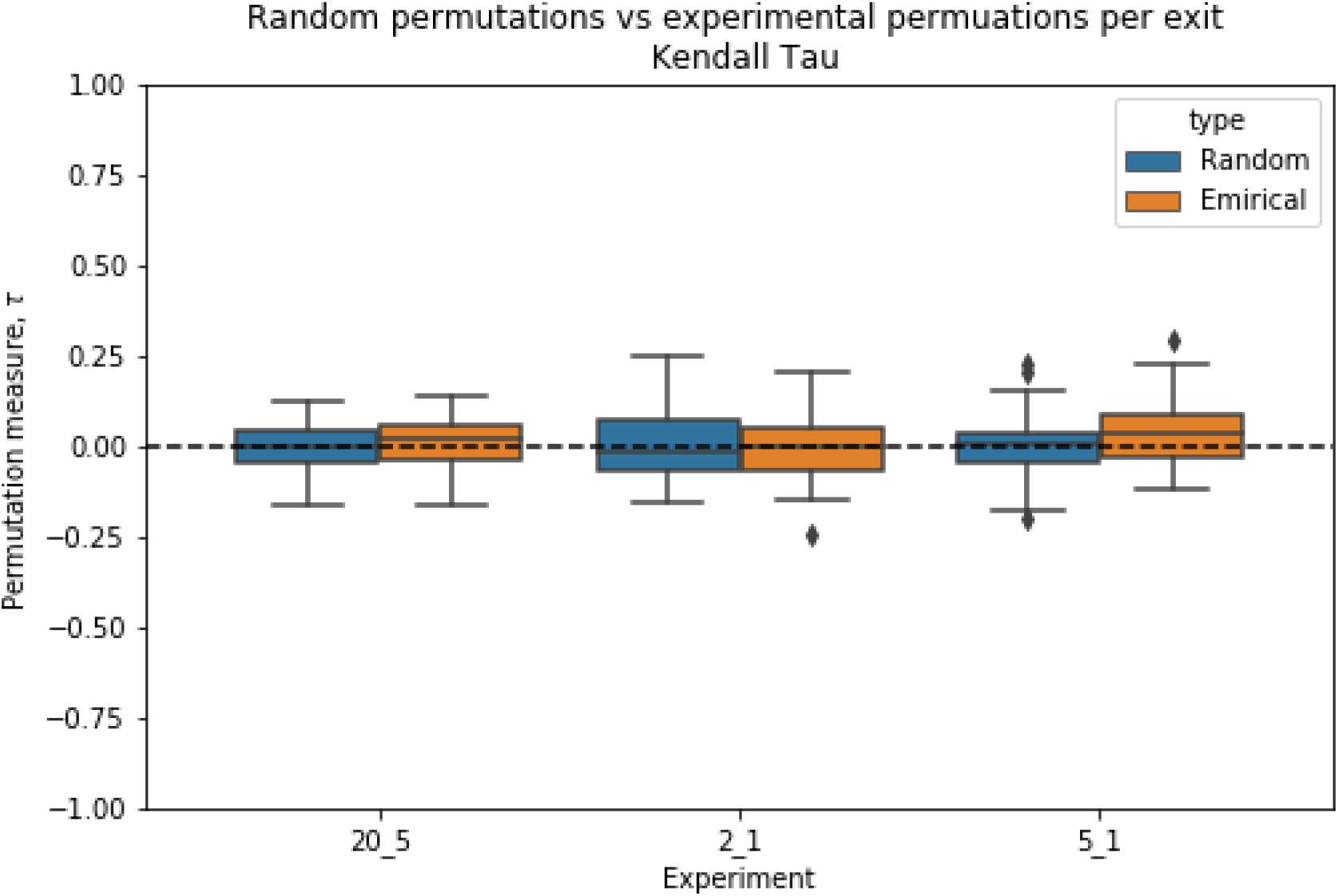
Distributions of the Kendall *τ* rank correlation coefficients for nest-ant movement in 3 experiments compared to those of fully random shuffling.

### Emergent linear relationship between foraging frequency and total colony hunger

In the main text, we have presented equations that explain the linear relationship between foraging frequency and total colony hunger for the extreme case, in which a forager walks only inward when her crop load exceeds the threshold, and only outward when it is below the threshold. Here we generalize these equations for the cases where the forager’s bias is set with partial probabilities to walk in each direction.

Let us describe the forager’s probabilistic walk as a random walk with a crop-load dependent bias *B*(*c*), where *c* is the forager’s current crop load, and *B*(*c*) is a set of 3 complementary fractions describing the probabilities to take one step inward, outward or to stay in the same cell, given *c*. In our case, a crop-load threshold *c*\* defines 2 biases: *B*(*c* > *c*\*) where the probability to step inward is greater than the probability to step outward, generating a net inward drift, and *B*(*c* ≤ *c*\*) where the probability to step outward is greater than the probability to step inward, generating a net outward drift. We will denote these biases *B_in_* and *B_out_*, respectively. For example, the biases used in the agent-based model are presented in Table 1, the first row corresponding to *B_out_* and the second to *B_in_*.

When compared to the extreme case presented in the main text, the main difference that this probabilistic walk introduces is that as the forager walks stochastically back and forth between the nest ants, she may interact multiple times with the same ants before switching her bias. For any biased random walker walking on an infinite line, the average number of times it steps on a specific position is a constant, the value of which depends on the value of the bias. This is due to the fact that a biased random walk is Markovian, such that the direction of the next step is independent of the walker’s position. Therefore, we can separately analyze the two phases of the forager’s walk in the nest: (1) when she enters the nest and drifts inwards with bias *B_in_* until she reaches her crop-load threshold, and (2) after she reaches the threshold and drifts back toward the nest entrance with bias *B_out_*. During each one of those phases, the average number of times the forager interacts with each ant is a different constant, which we denote *s*(*B_in_*) and *s*(*B_out_*), respectively. Note that this holds under the assumption that the forager is walking far enough from the nest boundaries. The average number of encounters with ants that are close to the boundaries may depend on their position, but in any case should stay quite constant during the entire course of the feeding process, and therefore we neglect this complication for the sake of our analysis.

Similarly to the extreme case presented in the main text, the nest-ant movement rule dictates that the forager interacts with a representative sample of the colony at each trip, such that the average state of the forager’s partners is equal to the colony state, *F*. However, for the probabilistic walk, this is true only for the first time the forager interacts with this partner. When repeatedly giving food to the same ants, their crop state gradually increases, affecting the amount of food they will receive in each successive interaction. Therefore, given the trophallaxis rule, by which the average amount of food delivered at each interaction is a fraction *α* of the receiver’s empty crop space, the total amount of food given to an ant who has been interacted with *s* times is, on average:

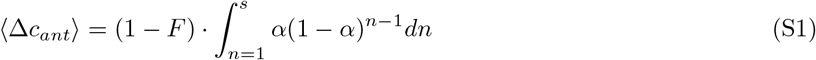

Now, the average position at which the forager reaches her threshold can be expressed as the average number of unique ants she interacts with before unloading 1-*c*\* of her crop. For the probabilistic case, this equals:

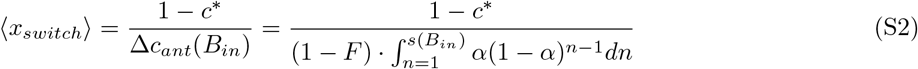

Since *s*(*B_in_*) and *α* are constants, 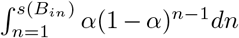 is also a constant, which we denote *η*, and obtain:

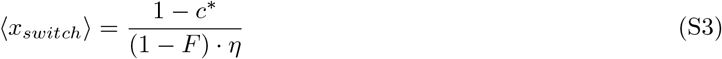

As in the extreme case, the average duration of a forager’s trip in the nest is the average time it takes her to reach 〈*x_switch_*〉 from the entrance plus the average time it takes her to reach the entrance back from *x_switch_*. For the probabilistic case, these may be expressed as 〈*x_switch_*〉 · *s*(*B_in_*) and 〈*x_switch_*〉 · *s*(*B_out_*), respectively. Therefore, the average time the forager spends in the nest before exiting is:

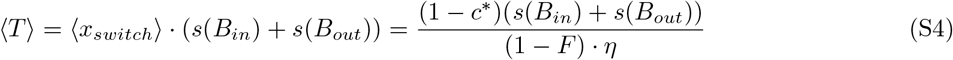

The exiting frequency is thus:

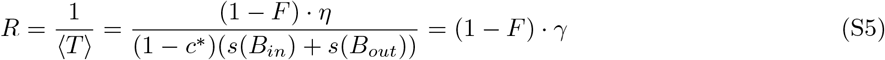

where 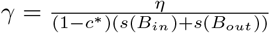 is a constant.

Hence, we get a foraging frequency *R* which is linear with colony hunger (1 – *F*) for the probabilistic case as well.

## Notes

### Competing Interest Statement

The authors have declared no competing interest.

